# Chromosome-level genome assembly of the Asian tramp snail *Bradybaena similaris* (Stylommatophora: Camaenidae)

**DOI:** 10.1101/2024.12.26.630445

**Authors:** Yasuto Ishii, Atsushi Toyoda, Alec Lewis, Angus Davison, Osamu Miura, Kazuki Kimura, Satoshi Chiba

## Abstract

While terrestrial land snails have long been subjected to evolutionary research, a lack of high-quality genomic resources has impeded recent progress. *Bradybaena* snails in particular have numerous intriguing traits that make them a good model for studying evolution, including shell pattern polymorphism and convergent evolution. They are also introduced and invasive across the world. In this study, we present a chromosome-level genome assembly of the Asian tramp snail *Bradybaena similaris* utilizing 88-fold Illumina short-read sequences, 125-fold Nanopore long-read sequences, 63-fold PacBio HiFi sequences, and 47-fold Hi-C sequences. The assembled genome of 2.18 Gb is anchored to 28 chromosomes, and exhibits high completeness (98.8% metazoan BUSCO completeness) and contiguity (N50 of 75.6 Mb). Additionally, we also obtained a high-quality transcriptome for annotation. This resource represents the first chromosome-level assembly for snails in the superfamily Helicoidea, which includes more than 5,000 species of terrestrial snails, and will facilitate genomic study in *Bradybaena* and, more broadly, in the superfamily Helicoidea.

**SIGNIFICANCE STATEMENT:** While the Helicoidea is the largest land snail superfamily, consisting of more than 5,000 species, many of interest to evolutionary studies, no chromosome-level assembly has been available for any species. Previously, the genus *Bradybaena* in the Helicoidea has been studied in the context of speciation, adaptation and invasive biology, and thereby has high potential for further research. In this study, we present a chromosome-level assembly and transcriptome of the Asian tramp snail *Bradybaena similaris*. These high-quality genomic resources will facilitate research on related species, and eventually enhance our understanding of many areas of evolutionary biology.

## INTRODUCTION

Terrestrial snails are relevant to many scientific fields. In the context of invasion biology, they are recognized as both agricultural and public health pests, leading to extensive research on their control (Hirano et al., 2020; Yonow et al., 2023). Unfortunately, in conservation biology terrestrial snails are recognized as holding a position of significant risk (Chiba and Cowie, 2016), representing the animal group with the most extinctions since the 16th century, with many species now requiring conservation efforts (Lydeard et al., 2016). Terrestrial snails also often serve as indicators of environmental pollution and so have been used in the study of anthropogenic impacts on the environment (Baroudi et al., 2020).

Terrestrial snails are a key group also in evolutionary biology (Chiba and Cowie, 2016), especially species in the Helicoidea, the most speciose land snail superfamily, including more than 5,000 species (MolluscaBase, 2024, accessed on 4th September 2024). For example, the genera *Satsuma* and *Euhadra* have been studied with the potential to understand speciation, and also the origins of variation in left-right asymmetry (Ueshima and Asami, 2003; Davison et al., 2005; Hoso et al., 2010; Richards et al., 2017). Also, research on *Theba*, *Cepaea* and *Euhadra* has been performed for the evolution of shell color (Knigge et al., 2017; Ito et al., 2023; Chowdhury et al., 2024). For another instance, in studies of behavioral evolution, *Cornu* and *Euhadra* have been major subjects of research on sexual conflict and mating behavior (Koene, 2006; Kimura et al., 2013).

Despite these advantages as excellent model systems, genomic research has been scarce in terrestrial snails, partly due to a lack of high-quality genome assemblies (Davison and Neiman, 2021; Linscott et al., 2022). As of now, genome assemblies are available for only eight land snail and slug species (Table S2). Among these, only the five species of *Achatina*, *Arion*, *Megaustenia* and *Meghimatium* have chromosome-level assemblies (Guo et al., 2019; Liu et al., 2021; Chen et al., 2022; Chetruengchai et al., 2024; Sun et al., 2024). Only two genome assembly has been published for Helicoidea (*Candidula*: Chueca et al., 2021; *Cepaea*: Saenko et al., 2021), and no chromosome-level assembly has been obtained.

*Bradybaena* snails in the Helicoidea are particularly intriguing in the context of speciation and adaptation, where similar shell morphologies have evolved independently (Hirano et al., 2014; 2019). As shell morphology is a key trait for assortative mating (Kimura et al. 2015), then this may lead to parallel speciation. In the context of adaptation, the shell color and band pattern of *Bradybaena* snails is also important. *Bradybaena* snails have polymorphism in shell color and band pattern, with the variation likely maintained through balancing selection (Komai and Emura, 1955). Each phenotype is determined by a single locus in the form of a supergene (Komai and Emura, 1955), a concept that has recently received renewed attention (Berdan et al., 2022). Also, *Bradybaena* snails are also highly invasive around the world. *Bradybaena similaris*, native to East and Southeast Asia, has been introduced to every continent except Antarctica (Serniotti et al., 2019). This species not only damages crop but also hosts parasites, some of which can cause diseases in humans and livestock (Serniotti et al., 2019). Genomic resources could support the further understanding of evolutionary innovations that make these invasive species, and by facilitating the development of control strategies (North et al., 2021).

Here, we represent a chromosome-level genome assembly of *B. similaris*. Using high-coverage Nanopore long-reads as well as PacBio HiFi reads, we successfully assembled a high-quality genome. Additionally, we obtained a high-quality transcriptome using newly sequenced RNA-seq data for annotation. The value in these genomic resources is their potential to facilitate research on *Bradybaena* snails, and more generally helicoidean snails, that can greatly benefit our understanding of multiple topics within evolutionary biology.

## RESULTS AND DISCUSSION

Sequencing yielded 192 Gb Illumina paired short-read sequences, 272 Gb Nanopore long-read sequences, 136 Gb PacBio HiFi sequences, and 101 Gb Hi-C library sequences (Table S1). Genome size and heterozygosity estimated by a k-mer-based method were 1,774,934,765 bp and 1.61%, respectively (Fig. 1B; Ranallo-Benavidez et al., 2020). An initial assembly generated by hifiasm (Cheng et al., 2022) had a size of 2,289,349,004 bp, consisting of 279 contigs, with N50 of 17,829,327 bp. This was then scaffolded using Hi-C reads, producing a final assembly of 2,179,927,975 bp, comprising 82 scaffolds (Table 1). Of these, 54 scaffolds were anchored onto 28 megascaffolds (Fig. 1C), which corresponds to the haploid chromosome number of the species (Inaba, 1959). The assembled genome size was larger than the genome size estimated using the k-mer method, a discrepancy that has been observed in other large-genome organisms, and is likely due to a high proportion of repeats (Pfenninger et al., 2022; Miura et al., 2023). The assembled genome has a higher contiguity and completeness than that of the other terrestrial snails (Table S2).

**Fig. 1.**
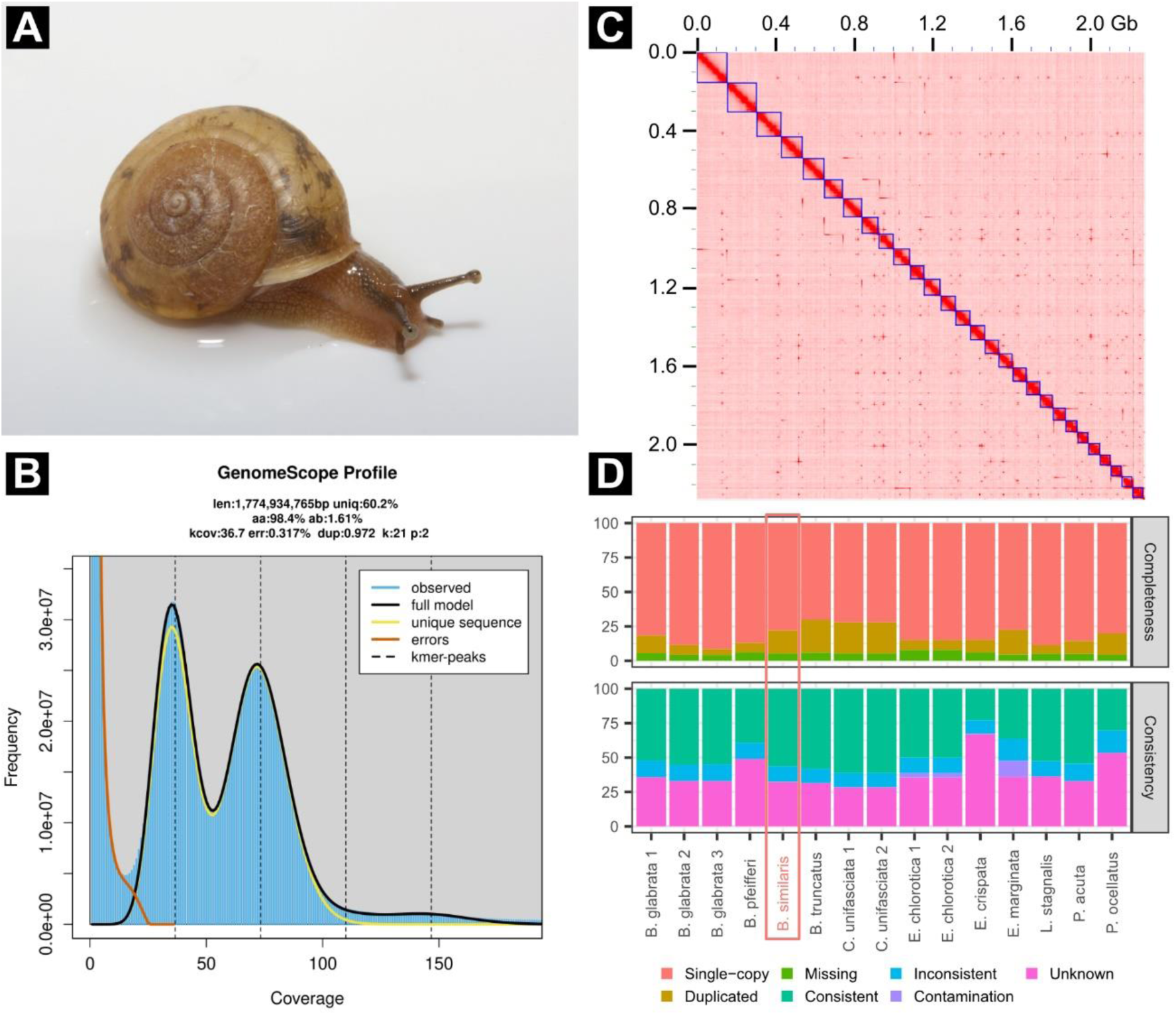
Overview of the results. (A) Photograph of *Bradybaena similaris*. (B) Genomescope k-mer profile plot for *B. similaris*, displaying observed k-mer frequency (blue bar), full model fitted by Genomescope (black line). (C) Hi-C contact map for *B. similaris* with chromosomal scaffolds (blue boxes). Darker red indicates higher contact density. (D) Comparison of OMark statistics for Panpulmonata snails. All data was sourced from OMark web server. The shown species and accession numbers (in parentheses) are following: *Biomphalaria glabrata* (BglaB1, UP001165740 and GCF_947242115.1), *Biomphalaria pfeifferi* (GCA_030265305.1), *Bradybaena similaris* (this study), *Bulinus truncatus* (GCA_021962125.1), *Candidula unifasciata* (UP000678393 and GCA_905116865.2), *Elysia chlorotica* (UP000271974 and GCA_003991915.1), *Elysia crispate* (GCA_033675545.1), *Elysia marginata* (GCA_019649035.1), *Lymnaea stagnalis* (GCA_964033795.1), *Physella acuta* (GCF_028476545.1), and *Plakobranchus ocellatus* (GCA_019648995.1).

**Table 1.**
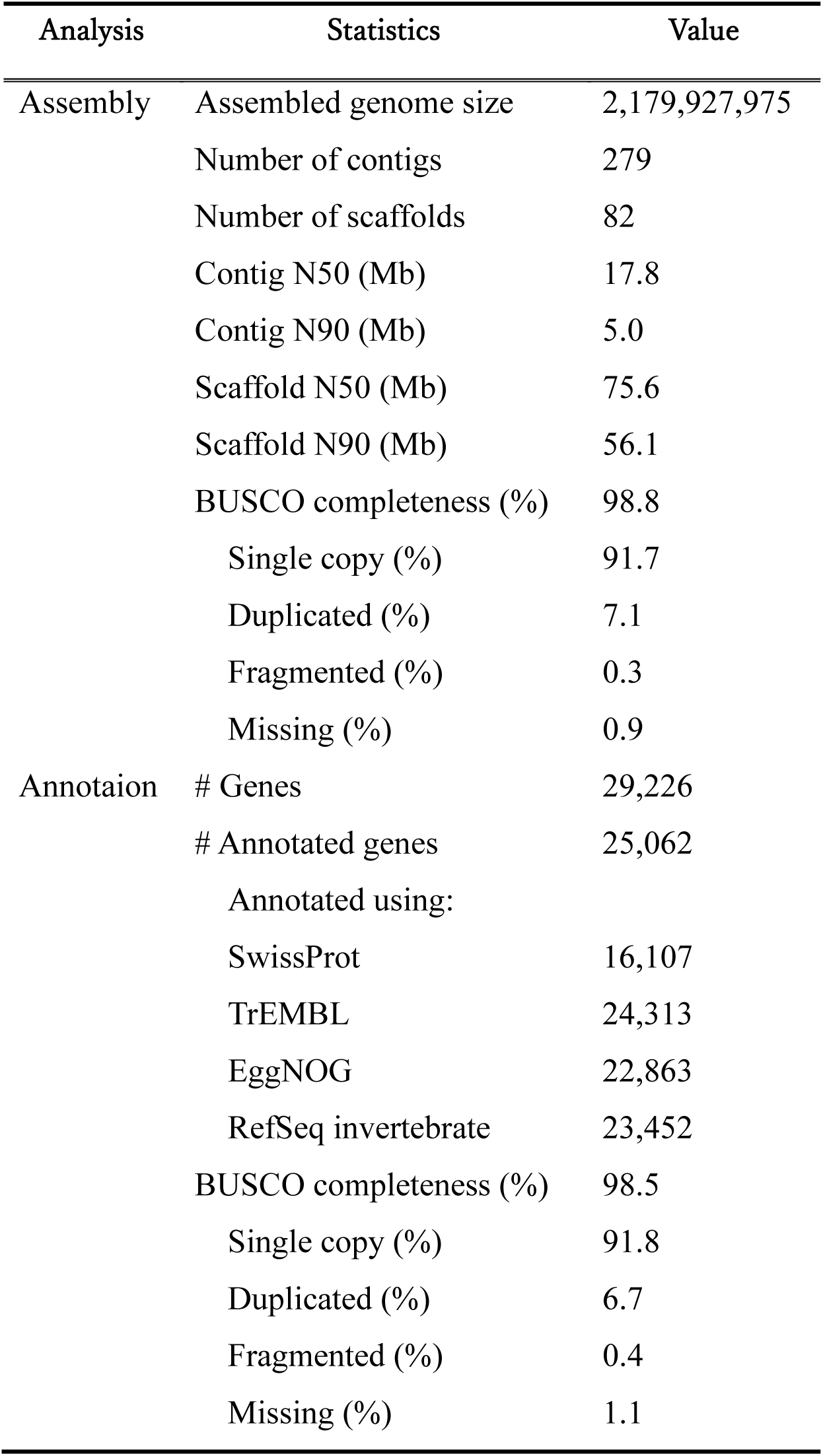
The statistics for the genome assembly and annotation for *Bradybaena similaris*, after removing contaminants.

The proportion of repeat content was estimated to be 61.28% (1.40 Gb), with long interspersed nuclear elements (LINEs) constituting a significant portion of the repeat content (19.51%; details are shown in Table S3). The proportion of LINEs is smaller than those of some terrestrial snails and slugs: *Candidula unifasciata*, 25.0% (Chueca et al., 2021); *Cepaea nemoralis*, 30.5% (Saenko et al., 2021); *Meghimatium bilineatum*, 34.1% (Sun et al., 2024).

The genome was annotated using BRAKER3 (Gabriel et al., 2024), with 87.91 Gb RNA-seq data and OrthoDB 11 (Kuznetsov et al., 2023). The number of predicted genes was 29,227. Functional annotation was performed for the predicted genes, and contaminants were removed using EnTAP (Hart et al., 2020). Any gene that was identified as putatively of bacterial origin was regarded as a contaminant and removed. Using four protein sequence databases as a reference (SwissProt, TrEMBL, EggNOG, RefSeq invertebrate), a total of 25,062 genes were functionally annotated (Table 1). Among the annotated genes, 16,631 genes had Gene Ontology terms contained in EggNOG (Table 1). Only one gene was identified as a putative contaminant, so 29,226 genes were finally retained. The number of genes in the final transcriptome is comparable to those of other land snails (Table S3).

Almost all metazoan core genes were found in the assembly and the final transcriptome (98.8% and 98.5%, respectively; Table 1). The quality of the transcriptome was also assessed with OMArk (Nevers et al., 2024), which quantifies not only the completeness as BUSCO does (i.e., measuring how many expected genes are present in the genome), but also the consistency (i.e., the proportion of protein sequences correctly assigned to known gene families within the same lineage: Nevers et al., 2024). According to OMArk, 56.39% of the transcriptome was classified as "consistent," 11.24% as "inconsistent," and 32.37% as "unknown". No gene was classified as "contaminant". Although the relatively high proportion of inconsistent genes might suggest that the genome annotation can be improved, the quality was higher than that of most other Panpulmonata proteomes (i.e., higher "consistent" rate and lower "inconsistent" or " unknown" rate; Fig. 1D). The lack of model species in related taxa (Davison and Neiman, 2021) may lead the relatively high proportion of inconsistent and unknown genes may be due to the lack of model species in related taxa, as similar trends have been observed in other Panpulmonata proteomes (Fig. 1D).

## CONCLUSIONS

We presented a chromosome-level genome assembly for *B. similaris* utilizing Illumina paired-read sequences, Nanopore long-read sequences, HiFi long-read sequences and Hi-C sequences. The assembled genome was 2.18 Gb in size, with N50 of 75.6 Mb and 98.8% metazoan BUSCO completeness. This quality is much higher than that of the other published terrestrial snail genomes. The estimated transcriptome was also high-quality, containing 98.5% metazoan BUSCO. The high-quality genome resources would accelerate the genomic study of terrestrial snails, particularly *Bradybaena* snails.

## MATERIALS AND METHODS

### Sampling, DNA extraction and sequencing

Three adult *B. similaris* individuals were collected for genome sequencing in Sendai, Miyagi, Japan (38.259661N 140.861679E) in September 2023 (Fig. 1A; Table S1). All snail shells were pale-colored and lacked bands. These three individuals were separately used for short-read sequencing, long-read sequencing, and Hi-C sequencing (Table S1). DNA was extracted from a piece of foot using Nanobind Tissue Big DNA Kit (PacBio, US) for short-read sequencing and QIAGEN Genomic-tip 100/G (QIAGEN, Germany) for long-read sequencing.

An Illumina short-read library was prepared with a TruSeq DNA PCR-Free Library Prep Kit (Illumina, US). Two runs of 150 bp paired-end sequencing was then performed on a NovaSeq 6000 (Illumina) using the NovaSeq 6000 SP Reagent Kit v1.5 (Illumina). The Nanopore reads were obtained using a PromethION, a Ligation Sequencing Kit V14 and a R10.4.1 flowcell (Nanopore, UK). The PacBio library was prepared using a SMRTbell prep kit 3.0 and sequenced on both Sequel II and Sequel IIe platforms, using a Sequel II SMRT Cell 8M along with a Sequel II Binding Kit 3.2 and Sequel II Sequencing Kit 2.0 (PacBio). These library preparations and sequencings were performed at the National Institute of Genetics Japan (NIG, Japan). An Hi-C library was prepared using Proximo Hi-C kit (Animal) following the manufacture’s instruction. Sequencing was conducted on a DNBseq-g400 (MGI, China) at BGI.

RNA-seq was performed for annotation. An additional individual was collected at Sendai, Miyagi, Japan (38.259550N 140.850820E; sample ID: MNKS 6468). Tissue of dart sac was preserved in RNAlater (QIAGEN) until extraction. Total RNA was extracted using ISOGEN (NIPPON GENE, Japan). Library preparation and sequencing was performed at BGI. DNBseq-g400 was utilized for 150 bp paired-end sequencing.

### De novo genome assembly

*De novo* assembly was initially performed with Hifisam 0.19.5-r587 (Cheng et al., 2022) using Nanopore as well as HiFi reads. Haplotigs were purged by purge_dups 1.2.5 (Guan et al., 2020), applying the "-e" option in the get_seqs module. The Hifiasm assembly was then used as input for Hi-C scaffolding. Hi-C reads were mapped onto the assembly using bwa 0.7.17-r1188 (Li et al., 2013) and samtools 1.17 (Danecek et al, 2021). Then, the mapped reads were used for Hi-C scaffolding using YaHS 1.2 (Zhou et al., 2023). The contact map was visualized, mis-joined scaffolds were manually split and the final assembly was generated using Juicer 1.2 (Durand et al., 2016), Juicer tools 1.9.9, and JuiceBox 2.15.0.0 (Robinson et al., 2018).

Genome features, including genome size, heterozygosity and repetitiveness were estimated using genomescope 2.0 (Ranallo-Benavidez et al., 2020). The canonical k-mer distribution was produced from the short-read data with KMC 3.1.1 (Kokot et al., 2017) with k-mer length and maximum coverage set as 21 and 10,000, respectively.

### Genome annotation

Prior to genome annotation, the repeat content was masked. RepeatModeler 2.0.4 (Flynn et al., 2020) was used to build a species-specific repeat content library. Using this library, the repeat content was masked using RepeatMasker 4.1.5 (http://www.repeatmasker.org). Both software tools were implemented in TETools 1.8 (https://github.com/Dfam-consortium/TETools).

Gene prediction was performed for the soft-masked sequence using BRAKER 3.0.8 (Gabriel et al., 2024). RNA-seq data (available at NCBI under accession numbers SRR8040510–SRR8040517, SRR6981555, and newly obtained data) and protein data (molluscan proteomes in OrthoDB 11; Kuznetsov et al., 2023) were used as reference data. The tissue for RNA-seq was sourced from whole body, embryo, digestive gland, and dart sac, among which dart sac data is newly obtained in this study. Functional annotation and contaminant removal for the predicted genes was carried out with EnTAP 1.3.0 (Hart et al., 2020). Four databases (SwissProt, TrEMBL, EggNOG, and RefSeq invertebrate) were referenced (O’Leary et al., 2016; Huerta-Cepas et al., 2019; UniProt Consortium, 2023). Contaminants derived from bacteria were discarded. Gene Ontology terms were assigned through EggNOG database.

The gene content completeness of the final assembly and the final transcriptome (predicted genes after contaminant removal) was assessed using BUSCO 5.7.1 (Manni et al., 2021), by searching against the metazoan core genes (metazoa_odb10). OMArk 0.3.0 (Nevers et al., 2024) was also used to quantify the quality of the annotation. To remove splicing variants, a transcript was extracted for each gene.

## ACKNOWLEDGEMENTS

We express our sincere gratitude to Mason Linscott, Yuki Kimura, Masanori Tatani, Christine Parent, Margrethe Johansen, Gemma Collins and Takahiro Hirano for technical support and advice. We also thank Sachiko Horie and Masayuki Maki for experimental assistance. Artificial intelligence was utilized for English editing. This work was supported by the Japan Society for the Promotion of Science (JSPS) KAKENHI (Grant Numbers 21H02556 and 22H04925 [PAGS], 22K06383 and 24K02094) and JSPS Summer Programme Fellowship.

## DATA AVAILABILITY

All raw data and assembly were deposited to DDBJ (Bioproject accession number: PRJDB16720). The accession numbers for each sequence are: DRR527377 for Illumina short-read sequences, DRR623704–623707 for PacBio HiFi sequences, DRR528210 for Nanopore long-read sequences, DRR624568 for Hi-C sequences and ### for RNA-seq sequences. [The RNA-seq data, genome assembly and transcriptome are about submitting, but unfortunately the DDBJ submission system is not available now, so the submission is still ongoing. All submissions are going to be completed until the end of the review process. All the data will be disclosed immediately after the manuscript is accepted].

