## Supplementary files for "Chromosome-level genome assembly of the Asian tramp snail *Bradybaena similaris* (Stylommatophora: Camaenidae)"

Genome resources

Short running title: **Chromosome-level genome assembly of *Bradybaena similaris***

Yasuto Ishii [1]*, Atsushi Toyoda [2], Alec Lewis [3], Angus Davison [3], Osamu Miura [4], Kazuki Kimura [1,5], Satoshi Chiba [1,5]

[1] Graduate School of Life Sciences, Tohoku University, Sendai, Japan

[2] Advanced Genomics Center, National Institute of Genetics, Mishima, Japan

[3] School of Life Sciences, University of Nottingham, Nottingham, UK

[4] Faculty of Agriculture and Marine Science, Department of Marine Resource Science, Marine Biological Chemistry Course, Kochi University, Kochi, Japan

[5] Center for Northeast Asian Studies, Tohoku University, Miyagi, Japan

### TABLES AND FIGURES (WITH CAPTIONS)

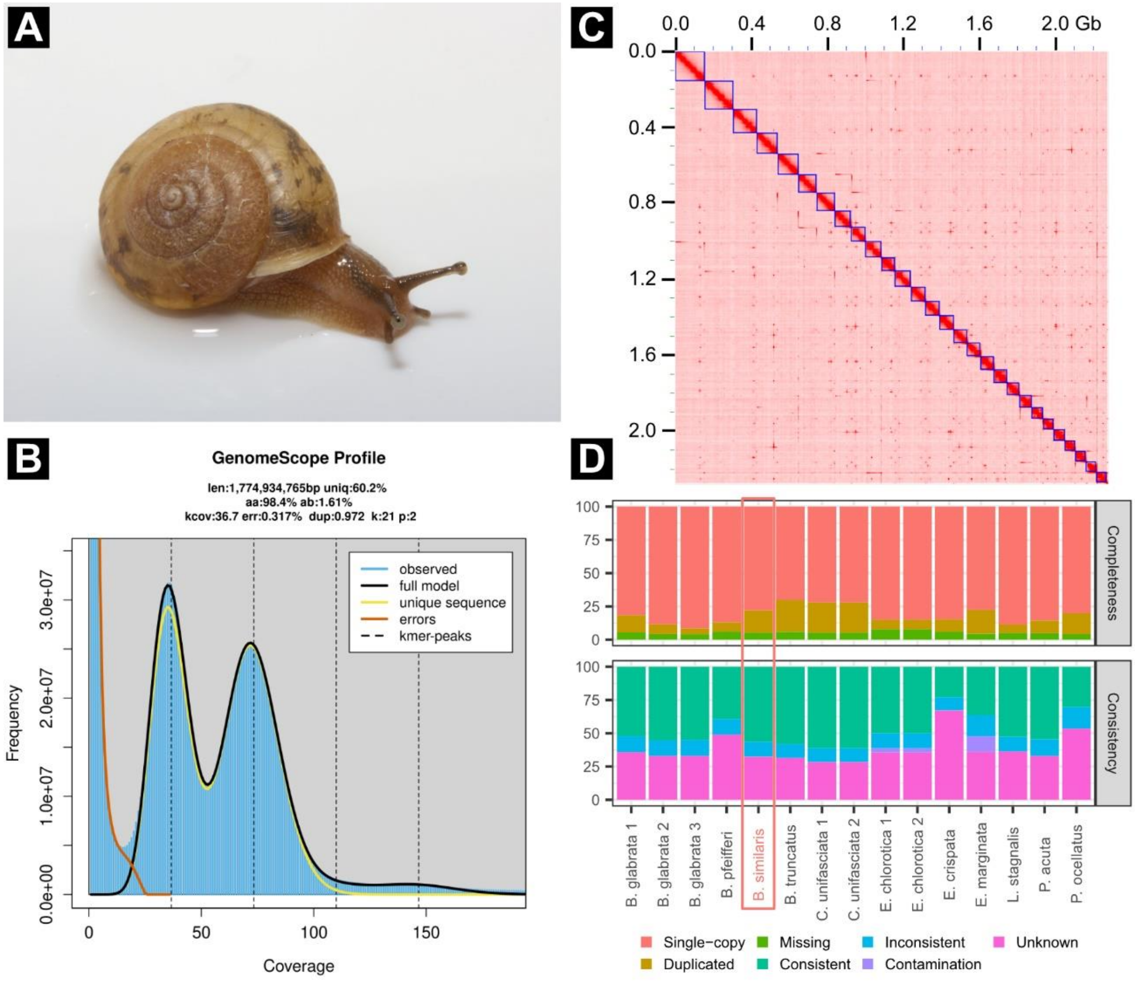


**Fig. 1. Overview of the results.** (A) Photograph of *Bradybaena similaris*. (B) Genomescope k-mer profile plot for *B. similaris*, displaying observed k-mer frequency (blue bar), full model fitted by Genomescope (black line). (C) Hi-C contact map for *B. similaris* with chromosomal scaffolds (blue boxes). Darker red indicates higher contact density. (D) Comparison of OMark statistics for Panpulmonata snails. All data was sourced from OMark web server. The shown species and accession numbers (in parentheses) are following: *Biomphalaria glabrata* (BglaB1, UP001165740 and GCF_947242115.1), *Biomphalaria pfeifferi*

(GCA_030265305.1), *Bradybaena similaris* (this study), *Bulinus truncatus* (GCA_021962125.1), *Candidula unifasciata* (UP000678393 and GCA_905116865.2), *Elysia chlorotica* (UP000271974 and GCA_003991915.1), *Elysia crispate* (GCA_033675545.1), *Elysia marginata* (GCA_019649035.1), *Lymnaea stagnalis* (GCA_964033795.1), *Physella acuta* (GCF_028476545.1), and *Plakobranchus ocellatus* (GCA_019648995.1).

Table 1. The statistics for the genome assembly and annotation for *Bradybaena similaris*, after removing contaminants.

| Analysis | Statistics | Value |
|---|---|---|
| Assembly | Assembled genome size | 2,179,927,975 |
| | Number of contigs | 279 |
| | Number of scaffolds | 82 |
| | Contig N50 (Mb) | 17.8 |
| | Contig N90 (Mb) | 5.0 |
| | Scaffold N50 (Mb) | 75.6 |
| | Scaffold N90 (Mb) | 56.1 |
| | BUSCO completeness (%) | 98.8 |
| | Single copy (%) | 91.7 |
| | Duplicated (%) | 7.1 |
| | Fragmented (%) | 0.3 |
| | Missing (%) | 0.9 |
| Annotaion | # Genes | 29,226 |
| | # Annotated genes | 25,062 |
| | Annotated using: | |
| | SwissProt | 16,107 |
| | TrEMBL | 24,313 |
| | EggNOG | 22,863 |
| | RefSeq invertebrate | 23,452 |
| | BUSCO completeness (%) | 98.5 |
| | Single copy (%) | 91.8 |
| | Duplicated (%) | 6.7 |
| | Fragmented (%) | 0.4 |
| | Missing (%) | 1.1 |
